# A Genetic Platform for Functionally Profiling Odorant Receptors Ex Vivo Using Olfactory Cilia

**DOI:** 10.1101/2021.09.28.462154

**Authors:** Masayo Omura, Yukie Takabatake, Eugene Lempert, Sigi Benjamin-Hong, Charlotte D’Hulst, Paul Feinstein

## Abstract

The molecular basis for odor perception in humans remains an enigma as odorant receptors (ORs) are notoriously difficult to study outside of their native environment. Efforts toward OR expression and functional profiling have been met with limited success due to poor efficiency of in vitro cell surface expression. Olfactory cilia are known to contain all components of the olfactory signal transduction machinery and can be placed into an ex vivo well-plate assay to rapidly measure robust, reproducible odor-specific responses. We now describe the isolation of cilia from mouse nose bioreactors for two human ORs, OR1A1 and OR5AN1, and reveal 10-100 fold more sensitivity compared to previously reported assays. A single animal can produce up to 4,000 384-well assay wells, and isolated olfactory cilia can be stored frozen and thus preserved. This pipeline offers a sensitive, highly scalable ex vivo odor screening platform that opens the door for decoding human olfaction.

**One Sentence Summary:** olfactory-cilia based odor screening platform.

## Introduction

Decoding the sense of smell has remained elusive for the last 30 years. General odors are recognized by olfactory or odorant receptors (ORs) and amines are detected by trace amine-associated receptors (TAARs): both receptor groups are seven transmembrane (7TM) receptors belonging to the G-protein Coupled Receptor (GPCR) superfamily. Mice express over 1000 intact OR genes and 14 TAAR genes while the human OR repertoire is composed of ∼400 intact genes and five TAAR genes (1-3). It has been exceedingly difficult to identify odors that activate ORs and even if an odor can be identified in vitro, one has to validate the observation in vivo, which is rarely done (4-6). What is needed is a manner to produce an OR in an olfactory neuron and rapidly characterize it. Olfactory sensory neurons (OSNs) express a single allele of one OR gene at a time. In mice, this translates to a mere ∼0.1% of the total number of OSNs in the main olfactory epithelium (MOE) expressing the same OR, thereby posing a challenge for measuring the activity from a single OR.

We have been studying how to increase OR gene choice in olfactory neurons (7) and have shown we can dramatically increase the total number of OSNs expressing specific OR coding sequences. We previously reported a genetic approach allowing us to functionally characterize ORs in their native environment, the olfactory neurons within the nose. As such, instead of 10,000 expressing one mouse OR, we now have at least 500,000 OSNs expressing our OR transgene (7). Additionally, our approach expressing ORs in the mouse olfactory system allows us to determine for the first time whether specific human ORs are fully functional in also providing neuronal and axonal identity.

In the mammalian olfactory system, the entire odor binding capacity of a neuron exists within olfactory cilia, which protrude from the dendritic knobs of the OSNs into the mucus of the MOE lining the nasal cavity. ORs in cilia detect odors through a G protein-coupled signaling cascade (Fig.1). Upon odorant binding, the GPCR cascade activates type III adenylate cyclase (ACIII), resulting in a rapid rise in the second messenger cyclic AMP (cAMP), which subsequently binds and opens cyclic-nucleotide gated channels to promote calcium influx. The initiation of the odor transduction pathway and its key enzymatic components are exclusively localized to olfactory cilia (8-12). Isolated olfactory cilia contain the full OR signaling protein machinery (13) and can be utilized ex vivo to measure odor-induced OR activation with an adenylyl cyclase assay (14).

**Fig. 1.**
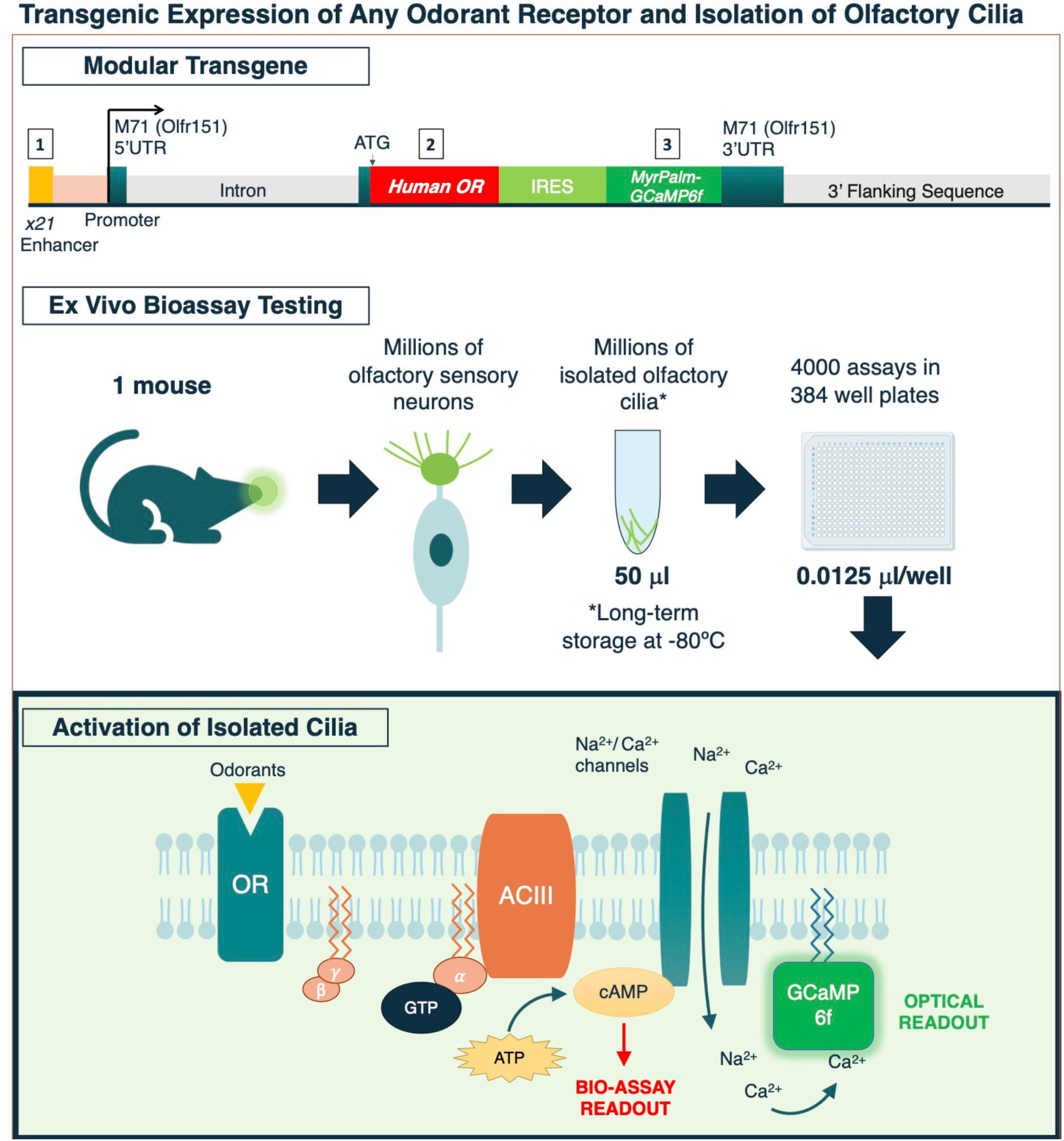
Ex vivo platform pipeline. Human Odorant receptors (ORs) are expressed using a modular transgenic construct. A high probability gene choice enhancer (5×21 or 9×21) sequence in a mouse OR gene, M71 (Olfr151) genomic backbone allows for the selective expression of the human OR together with a membrane targeted fluorescent reporter such as IRES MyrPalm-GCaMP6f. Transgenic mice express the OR and reporter in their olfactory epithelium, in which the olfactory cilia are extracted for use in an ex vivo bioassay. A biochemical assay measures cyclic AMP accumulation while a fluorescent readout measuring calcium influx is also possible.

While previous studies have used isolated cilia to measure OR activation by applying single odors or an odor mixture, this method alone cannot be used to pinpoint the activation of individual ORs in response to odorants (15, 16). Because each OR is found in approximately 0.1% of all OSNs, there are not enough cilia of a single OR type to define receptor-specific response profiles. Odor induced signals generated by all other ORs further complicate measuring signal output from a single OR type.

To circumvent these issues, we are taking a genetic approach to express human ORs in a large fraction of OSNs. Previously, we generated transgenes that increase the total number of OSNs expressing two control ORs, mouse M71 (Olfr151) and human OR1A1 by at least 250-fold (7). In this paper, we show that this dramatic overrepresentation provides the resolution necessary to identify odor responses unique to the over-represented OR. The expressed OR is easily determined as functional and physiologically relevant by examining the formation of green fluorescent glomeruli by tagging with GCaMP6f. A fundamental advantage of cilia isolation for ex vivo analysis over heterologous expression systems is they can be aliquoted and stored for prolonged periods of time (15-17). These cilia originate in transgenic animals that highly express the OR of interest. This is a clear advantage over heterologous systems that don’t include native binding and accessory factors in the membrane which are significant for proper OR signaling. Furthermore, we show that a green fluorescent sensor such as GCaMP6f (13, 18) can be targeted to olfactory cilia in transgenic lines, leaving open the possibility of measuring odor signaling optically within cilia (19, 20).

Our platform can generate an OR of our choosing to be expressed in a large number of OSNs such that cilia can be isolated and characterized with odors or ligands in a quick bench-top bioassay. The data reveal robust and reproducible nanomolar responses in liquid phase delivery of odorants. Vapor phase activation of ORs shows equivalent activation profiles as liquid delivery. OR activation is sensitive, specific and selective in a concentration dependent manner. This cilia-based platform finally opens a door for odor characterization of ORs including the entire repertoire of human ORs.

## Results

### Expression of Human Odorant Receptors

In order to functionally characterize ORs in cilia, we developed a strategy to 1) increase the number of OSNs expressing a given OR, and 2) isolate cilia for functional analysis.

We employed two well-characterized human ORs: OR1A1 and OR5AN1 (21-26). As we showed in our previous work (7), our platform uses a transgene that has three modules that can be readily modified: 1) the 5’end where either five or nine repeats of a gene choice enhancer (x21, ACATAACTTTTTAATGAGTCT) are used to boost the representation of a specific OR; 2) the CDS of Human OR, where any OR CDS can be inserted; and 3) a bicistronic reporter (IRES-Reporter) inserted 3’ of the CDS (Fig.1). To label all membranes in an OSN including cilia, we fused a previously characterized membrane targeting peptide tag (Myristylation/Palmitoylation-MyrPalm) to the green fluorescent reporter GCaMP6f (20). We generated two transgenic constructs: *OR1A1-IRES-MyrPalm-GCaMP6f* mice (OR1A1) and *OR5AN1-IRES-MyrPalm-GCaMP6f* mice (OR5AN1), both driven using a 9×21 enhancer. In 2/2 OR5AN1 and 1/1 OR1A1 transgenic mouse lines, we observe many more OSNs expressing the transgene in wholemount analysis of the epithelium and glomerular projections than the Olfr151 gene-targeted OR. In addition, MyrPalm-GCaMP6f robustly labels membranes of axons as they project to the olfactory bulbs and form large glomeruli (Fig.2). Notably, the OR5AN1 line has a greater number of projections in the Class I domain of the bulbs. Moreover, en face images of olfactory epithelium reveals brightly fluorescent labeled ciliary membranes (Fig.2) consistent with the MyrPalm lipid modification.

**Fig. 2.**
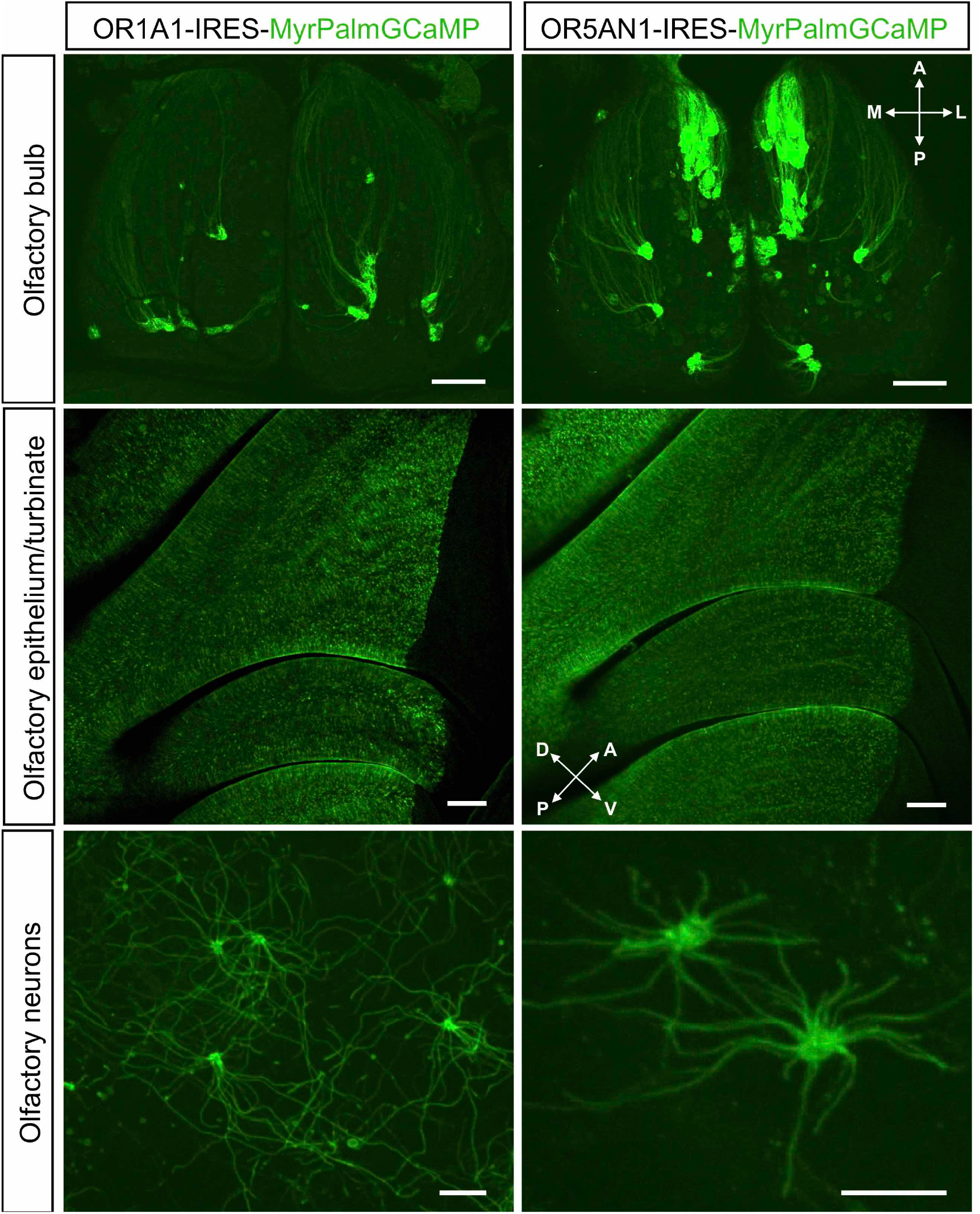
Myrpalm-GCaMP6f expressions in OR1A1 and OR5AN1 transgenic animals. The upper panels show the dorsal view of the olfactory bulbs of OR1A1-IRES-Myrpalm-GCaMP6f transgenic animals (left), and OR5AN1-IRES-Myrpalm-GCaMP6f (right), scalebar 500 μM. The middle panels show medial wholemount views of the main olfactory epithelium and turbinates of OR1A1 (left) and OR5AN1 (right), scalebar 200 μM. The bottom panels show high magnification en face view of the GCaMP positive olfactory sensory neurons of OR1A1 and OR5AN1animals, left and right respectively. A, anterior; P, posterior; L, lateral; and M, medial; D, dorsal; V, ventral, scale bars 10 μM.

It is believed that the olfactory sheet in the mouse nose is between 8-10 million neurons. There may be enough flexibility in the epithelium for the OSNs expressing cells from our transgenic lines generate additional cells or would usurp the probability of choice of the OR genes. We set out to quantify if other OR expressing neurons were lost and to what degree. We applied RT-qPCR for five OR genes that are highly expressed in the dorsal epithelium from both Class I and Class II cell types as well as one OR gene expressed in the ventral epithelium (Fig.S1). Both OR5AN1 and OR1A1 transgenic lines revealed a reduction of endogenous OR expression for 5/6 OR genes. The OR5AN1 line also showed a two-fold reduction in the two Class I genes tested relative to the OR1A1 line. These data suggest that a substantial portion of the dorsal epithelium has been coopted by the OR1A1 and OR5AN1 expressing OSNs consistent with the expression of GCaMP positive cells in the epithelium and substantial axonal projections in the bulb (Fig. 2).

### Functional Activation of ORs in from rapidly Extracted Olfactory Cilia

Odorant receptor signaling is self-contained in the cilia where ORs, Gαolf, ACIII and cyclic nucleotide-gated channel are localized; isolated cilia can readily convert ATP into cAMP in an odor dependent manner (Fig.1)(15). To study OR response profiles, we developed a pipeline to isolate cilia (15-17) and measure specific odor activation ex vivo. Cilia preparations were taken from transgenic and control mice using modified deciliation protocols (15-17, 26). We confirmed that Golf and ACIII were present in our isolated cilia preps by Western blotting as previously shown (9, 11). It has previously been described that isolated cilia can readily convert ATP into cAMP in an odor dependent manner (Fig.1). To detect OR activation by odorants in our isolated cilia enriched in human OR (hOR bioextracts), we used a modified adenylyl cyclase assay (15) combined with a commercially available cAMP kit to measure accumulated cAMP molecules in isolated olfactory cilia. Isolated cilia from a single mouse were placed in 10 × 5 μl aliquots and frozen. Each odor activation experiment requires a minimum of 0.0125 ul per well of a 384-well plate, yielding up to 400 reactions per 5 μl aliquot. The aliquoted samples are stable at least for 6-16 months at -80 °C without losing of basal cAMP production (86.00% ±38.5%, mean ±SD, 8 biological replicates, a total of 45 data points) nor losing activation with ligands (ΔcAMP value upon MND stimulation 97.37% ±38.3%, mean ±SD).

To test for receptor-specific responses, we compared cAMP responses in cilia from transgenic and non-transgenic control preparations using known OR1A1 and OR5AN1 agonists. To normalize for variability across ciliary preparations, we initially measured the maximum cAMP response in each preparation using the adenylate cyclase activator forskolin (FSK). Baseline cAMP production with DMSO or ethanol solvents range at 0.01-0.03 μM cAMP. By contrast FSK activation yielded ∼10-fold higher amounts (9.95 ± 0.81 - 10.61 ±0.66, mean ±SD) of cAMP between 0.05 and 0.38 μM (Fig.3A). We noted a strong correlation between the baseline amounts of cAMP compared to the FSK-activated cAMP (R^2^ = 0.953, Fig.3B). Thus, the baseline value of cAMP, DMSO induced cAMP, (cAMP^DMSO^) was used to normalize across experiments. Thus, ligand induced cAMP (cAMP^ligand^) - (cAMP^DMSO^) divided by cAMP^DMSO^ provided the measure of activation for each cilia sample.

**Fig. 3.**
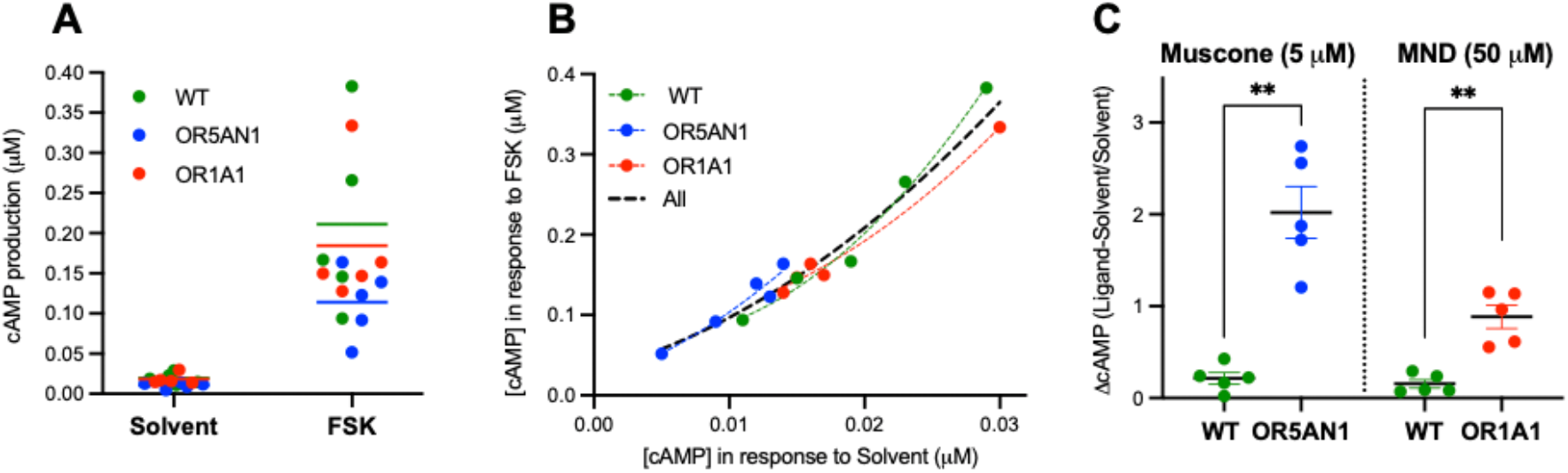
Ex vivo activation of human ORs with ligands. (**A**) Activation of wild type control (WT), OR5AN1 and OR1A1 hOR bioextracts with 5 µM forskolin (FSK), and the solvent, the bars represent mean and dots show datapoint of produced cAMP bars represent mean and dots show data points of produced cAMP by solvent and FSK, all groups are n = 5 biological replicates. (**B**) Correlation between cAMP in response to FSK (vertical axis) versus to solvent (horizontal axis). The dots show data points from 5 biological replicates with best fit nonlinear regression curves. (**C**) Activation of WT and OR5AN1 hOR bioextracts with 5 µM muscone, n = 5 biological replicates, and activation of WT and OR1A1 hOR bioextracts with 50 µM 3-methyl-2,4-nonanedione (MND), data represents activation values: mean of, ΔcAMP = (cAMP^ligand^ - cAMP^solvent^)/cAMP^solvent^ with ± SEM, superimposed data points, n = 5 biological replicates. Mann-Whitney test, **p < 0.01.

We tested for ligand response selectivity of OR5AN1 and OR1A1 to their reported ligands (5 μM muscone for OR5AN1 and 50 μM MND, 3-methyl-2,4-nonanedione, for OR1A1). OR5AN1 demonstrated a significant response to its published ligand muscone (25) at 5 µM (ΔcAMP = 2.02 ±0.028, mean ± SEM) compared to non-transgenic wild-type (WT) cilia (ΔcAMP = 0.21 ± 0.065, mean ± SEM) (Fig.3C, left). Similarly, OR1A1 responded to its published ligand MND (21) at 50 µM (ΔcAMP = 0.88 ± 0.065, mean ± SEM) (Fig.3C, right) compared to WT values cilia responses.

We further characterized OR5AN1 using a panel of 12 hedonically similar musk compounds of various chemical structures (Fig.S2). In agreement with recent reports (22, 23, 25, 27), OR5AN1 selectively responds to macrocyclic (compounds M1-M7), polycyclic (compound P1), and nitro musks (compounds N1-2) by demonstrating low micromolar to nanomolar sensitivity while showing a complete lack of response to alicyclic (compound A1-A2) musks (Fig.4 and S3). We demonstrate that the assay can discern affinity for various compounds based on 1) response intensity: the magnitude of the response to musks, and 2) response sensitivity: presence of response after serial dilutions of compounds.

**Fig. 4.**
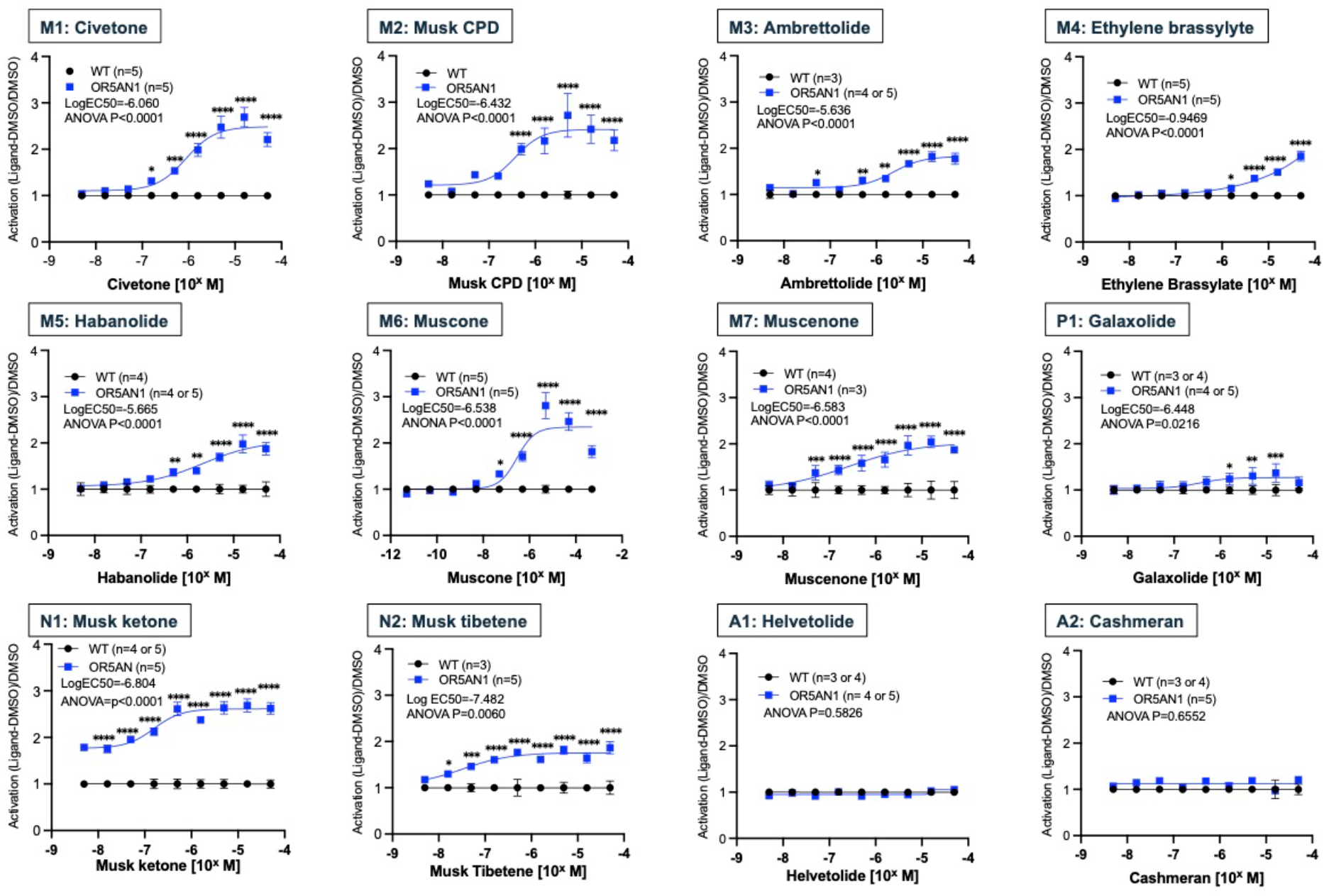
Ex vivo activation of OR5AN1 with 12 musk compounds. Activation of wild type control (WT) and OR5AN1 hOR bioextracts using a nine-point log dilution curve of 12 musk compounds, the black circles and blue squares, WT and OR5AN1 respectively, show mean of activation value (ΔcAMP = (cAMP^ligand^ - cAMP^solvent^)/cAMP^solvent^) corrected with its WT value=1 ± SEM. n=3-5 biological replicates as indicated in each graph. EC50 was calculated from the best fit nonlinear regression curve (four parameters) with the corrected activation values. Two-way ANOVA was performed, and p-values of the interaction were as indicated. Fisher’s LSD test was applied as post hoc comparisons, *p < 0.05, **p < 0.01, ***p < 0.001, ****p< 0.0001.

We next sought to define dose-response curves. However, high concentrations (≥ 5 µM) of certain musks had a deleterious effect on cAMP production, even in WT cilia (Fig.S3). This suggests interference with cAMP production perhaps from introducing high concentrations of hydrophobic compounds in the ciliary membrane. Notably, we observe activation to drop below 1 when administering higher concentrations, preventing a simple calculation of EC50s. Despite this drawback, we were still able to observe clear differential response to musks previously reported as weak or non-ligands of OR5AN1, ambrettolide (M3), ethylene brassylate (M4), habanolide (M5), and galaxolide (P1). Moreover, we controlled for the reduction in response by normalizing responses, shifting all WT to 1.0 and correcting corresponding Tg values (Fig.4). We find that musk tibetene has the lowest EC50 at -7.482 M, but not the strongest activation, by contrast, musk ketone has slightly low EC50s at -6.804 M, while having higher activations (Fig.4.).

### Selectivity, Specificity, Sensitivity, Efficacy, Reproducibility and Repeatability of OR responses

Given that odors normally enter the nose as volatile compounds present in the headspace, we next tested whether our ex vivo odor bioassay would be amenable to vapor phase delivery. To perform this series of experiments, we built a benchtop olfactometer with a custom Teflon manifold allowing delivery of up to 6 individual odors to 16 different cilia preps on a 96-well plate. We have observed little to no crosstalk from neighboring odor line chambers (Fig.S4) and can modulate concentration of delivered headspace by air dilution (Fig.S4). Delivery of headspace saturated with (-)-carvone (Saturated Vapor Pressure, SVP: 23.5 µM) elicited a response from the OR1A1 sample that was significantly above either WT or OR5AN1 responses (Fig.5A). Similarly, muscenone (SVP: 5.92 µM) differentially stimulated OR5AN1 compared to either WT or OR1A1 (Fig.5B). We observed that the odor delivery of compounds by vapor phase (Fig.5) is comparable to that of the liquid phase at the equivalent concentrations (Fig.4).

**Fig. 5.**
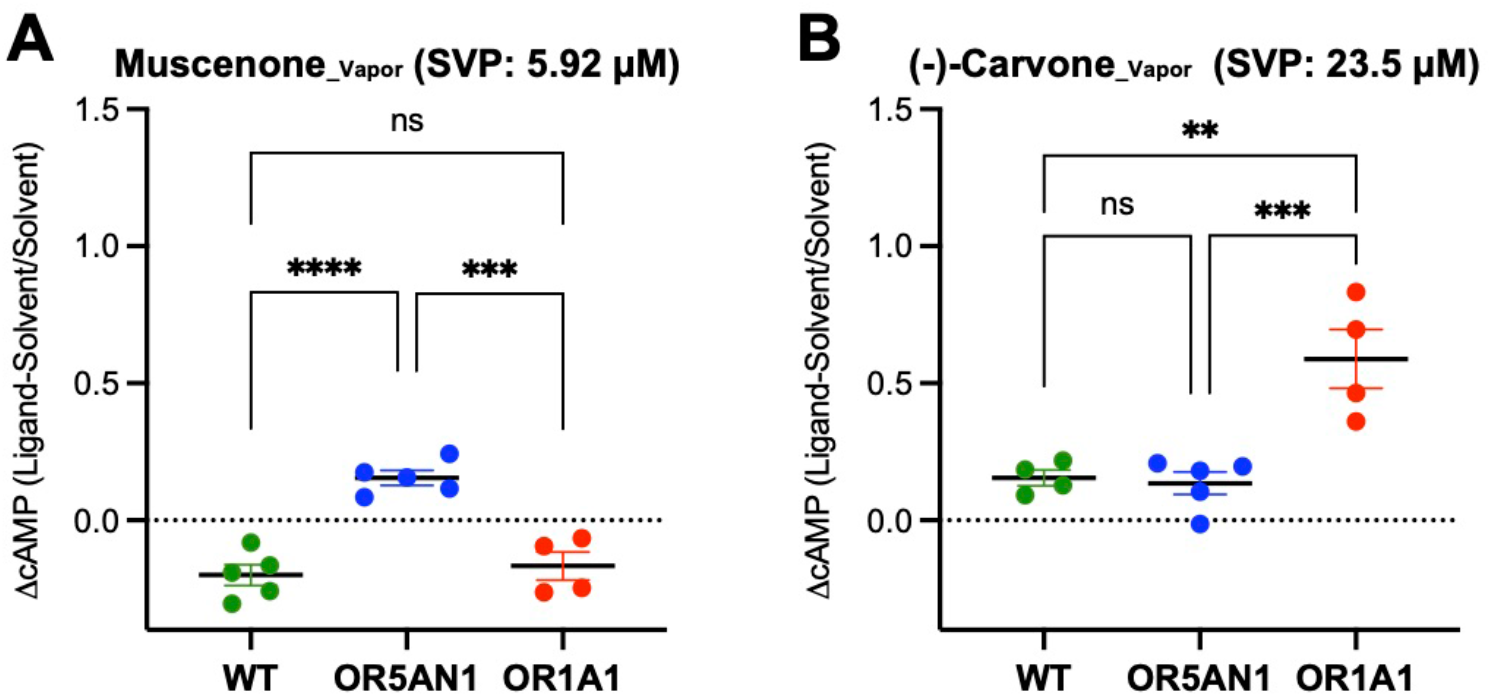
Headspace odor activation of OR1A1 and OR5AN1. (**A**) Activation, ΔcAMP = (cAMP^ligand^ - cAMP^solvent^)/cAMP^solvent^ of wild type control (WT), OR1A1 and OR5AN1 hOR bioextract using a vapor-phase delivery of Muscenone at saturated vapor pressure (SVP) concentration, 5.92 μM. WT; n = 5, OR5AN1; n = 5, OR1A1; n = 4 biological replicates, error bars show SEM. Ordinary One-way ANOVA with Fisher’s LSD test for each pair compared, ***p<0.001, ****p < 0.0001. (**B**) Activation of WT, OR1A1 and OR5AN1 hOR bioextract using a vapor-phase delivery of (-)-carvone at SVP concentration, 23.5 μM. WT; n = 4, OR5AN1; n = 5, OR1A1; n = 4 biological replicates, error bars show SEM. Ordinary One-way ANOVA with Fisher’s LSD test for each pair compared, **p < 0.01, ***p < 0.001.

MND, 3-methyl-2,4-nonanedione, is the highest affinity odor for OR1A1; two lower affinity ligands are the stereoisomers (-)-carvone and (+)-carvone (21, 24).We tested if the OR1A1 cilia would distinguish these stereoisomers. These lower affinity ligands showed differential responses in the 9×21-OR1A1-GCaMP6f cilia with (-)-carvone eliciting greater activation than (+)-carvone. However, the data were more easily quantified using our 5×21-OR1A1-Cherry line (7) which has about 5.06 ± 1.30 (mean ± SEM) times higher OR1A1 transcripts in the MOE than its in OR1A1-GCaMP animals (log2 Fold Change: 2.34 ±0.378 (SEM), OR1A1-Cherry; n=5, OR1A1-GCaMP; n=4). Consistent with differing OR1A1 transcript amounts, we find OR1A1-Cherry bioextracts showing 7.37 ± 1.24 times (mean ± SEM, n = 3, as a mean at the three concentrations, 5 μM to 500 μM) greater responses to MND than the OR1A1-GCaMP animals (Fig.S5A). In sum, OR1A1 ligand induced ΔcAMP values are higher for MND followed by (-)-carvone and lastly (+)-carvone (Fig.6A). If we normalize the fold activation for these three odors and take the average of 6 independent measures, MND is responding 3.5x over (-)-carvone, which in turn is responding 6.1x over (+)-carvone (Fig.6B).

**Fig. 6.**
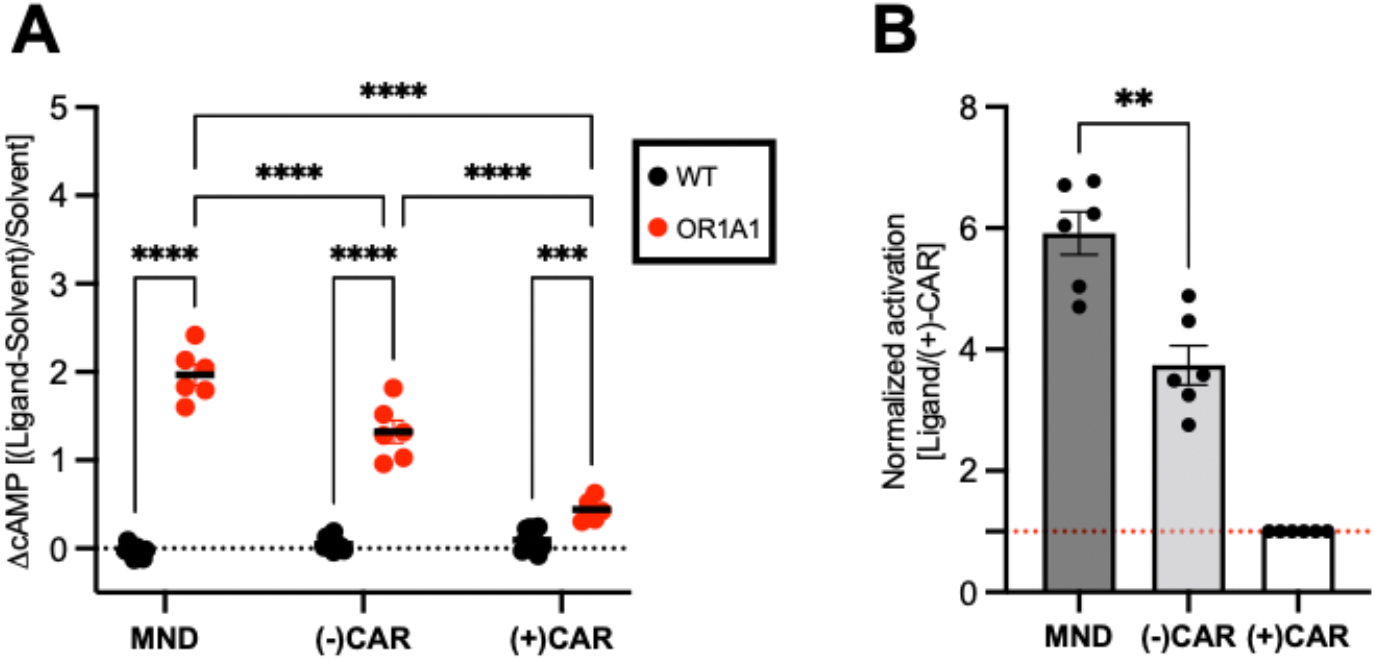
Comparison of ex vivo activation of OR1A1 with three known ligands. **(A)** Activation, ΔcAMP = (cAMP^ligand^ - cAMP^solvent^)/cAMP^solvent^) of wild type control (WT) and OR1A1 bioextracts with 3-methyl-2,4-nonanedione (MND), (-)-carvone [(-)-CAR], and (+)-carvone [(+)-CAR], shown with black dots and red dots, WT and OR1A1 respectively with the mean as black bars. WT; n = 10, OR1A1; n = 6. Two-way ANOVA was performed with Fisher’s LSD test was applied as post hoc comparisons, ***p = 0.0003, ****p < 0.0001. (**B**), ΔcAMP of MND and (-)-carvone were normalized with own ΔcAMP of (+)-carvone. The red dash line shows one. The right three columns show mean of the six OR1A1 data points represented as dots with SEM. Mann-Whitney test for normalized MND activation and (-)-CAR, **p = 0.0043.

## Discussion

### Heterologous expression systems

Heterologous expression systems have provided a wealth of pharmacological profiling for hundreds of GPCRs and the development of novel agonists and antagonists. These non-chemosensory GPCRs are typically characterized by robust plasma membrane expression in heterologous systems (28). However, nasal chemosensory GPCRs (Class I and II ORs as well as vomeronasal receptors) have exhibited poor plasma membrane expression in heterologous systems no matter the organism of origin (29).

It is estimated that only 10% have one identified ligand among the ∼400 intact human odorant receptors (30). Heterologous OR expression systems would lend themselves for high-throughput ligand characterization, but 30 years of effort has failed to decode how humans identify odors. To date, we still don’t understand why expression of the unmodified OR protein does not traffic to the plasma membrane outside of the olfactory neuron (31). Class A GPCRs do not have a signal peptide, but rather use the first transmembrane domain as a “signal domain” for insertion into the plasma membrane. In an effort to force plasma membrane expression in heterologous systems, many laboratories have added signal peptides to the amino-termini of these 7TM proteins or made chimeric GPCRs whereby the first and last transmembrane domains belong to non-chemosensory GPCRs (32). These modifications could disrupt the true odor profile of an OR. A promising line of research identified the putative GPCR cilia trafficking co-factors RTP1 and REEP1 based on functional screens looking for single pass proteins like odr-4. A third cofactor Ric8b was found in a similar manner (33). But, using both amino terminal tagging of ORs and cofactors, still produces very little protein on the plasma membrane compared to two canonical GPCRs beta2 adrenergic receptor or rhodopsin (34). Recent advances continue in heterologous expression of ORs using novel secretory signals such as the IL6-Halo-Tag system (35), which may prove valuable in identifying ligands for human ORs.

### Advantages of functional expression of cilia

Our approach obviates difficulties with heterologous expression by expressing ORs in native OSNs. This allows for proper OR localization to cilia plasma membrane as well as the natural environment for G-protein coupling. We show that comparing responses in WT cilia vs responses in cilia from transgenic overexpressing mice allows us to quantify odorant response properties of specific human ORs. Indeed, our calculated activation revealed odor-OR activities at nanomolar concentrations for high affinity ligands. The sensitivity of OR5AN1 for muscone is 40-100 fold better than heterologous expression assays with our EC50 at 0.29 μM (Fig.4) compared to 12.5-32.1 μM (22, 25) or 1000 fold more robust than another laboratory (36). Our OR1A1-Cherry mice with many more cells than the OR1A1-GCaMP6f mice has about 5 times higher OR1A1 transcripts expression and has higher sensitivity to ligands (Fig. S5A). The Cherry line has EC50 388 nM and a threshold of sensitivity to MND in the low nM range shown in Figure S5B, the data is in the same response range as the sensitive form of MND as shown by Geithe et. al., 2017 (21). Moreover, our assay can be used with any OR, even those that are resistant to functional expression in heterologous cells.

Another advantage of our assay is that it allows us to evaluate all required functions of ORs. In addition to binding odorants, ORs promote neuronal maturation and are required for proper axon guidance. Thus, for an OR to be used for ligand/odor detection, it must retain all three functions (37). In this context, if a full-length OR fails to do so, then we would consider it an intact pseudogene or non-functional gene. It is difficult to examine all three OR functions in humans. In fact, single cell sequencing of human OSNs has revealed around 140 out of ∼400 ORs with high amounts of expression suggesting that their neurons have matured (38). Our assay provides a method to determine full functionality of human ORs by expressing them under OR gene choice control mechanisms and assessing stable glomerular formation, as we have done for OR1A1 and OR5AN1. In this view, human ORs have not been fully characterized until their ability to support OSNs maturation and glomerular formation has been assessed.

Our approach is also scalable. Our highly represented ORs in mice provide enough material for maximum 4000 assays from one mouse to produce signals that are easily distinguished from non-transgenic (wildtype) signals when using a high affinity ligand. We show p-values at 0.001 for Transgenic OR over WT when responses were observed in the nanomolar range as shown in Fig.4.

### Odorant receptors versus chemical detection devices

Analytical chemistry characterizes chemicals by a subset of its features (light interaction, heat interaction, electric or magnetic field interactions). Olfaction is a measure of complex intermolecular forces (hydrogen or ionic bonding, ion-dipole or ion-induced dipole forces, van der Waals forces) between a chemical and a protein such that these intermolecular forces (ligand binding) have been optimized over evolutionary time to produce high affinity states that GPCR systems translate into intracellular signals in seconds (39). One molecule of a ligand can lead to the activity of hundreds of G-proteins each capable of stimulating adenylate cyclase activity leading to large numbers of cAMP molecules. The strength of GPCR activation as we show provides differential responses over a 2-log range and saturate over another 2-3 log range (40, 41). Taken together, ORs provide the signal amplification, specificity, efficacy that dwarfs any non-specific chemical detection device.

### Future directions for the platform

Our cilia preparations from the dorsal recess of the olfactory epithelium are not purely derived from one type of OR. Thus, the more Tg-OR derived cilia in a preparation, the more signal will be obtained as we see when comparing our Tg for 5×21-OR1A1-Cherry to 9×21-OR1A1-GCaMP6f (Fig. S5A). We recently characterized 5×21-OR1A1-Cherry (42) that reveals the dorsal epithelium is now expressing OR1A1 in 50% of the OSNs. Further increasing expression in the dorsal epithelium to 100% of the OSNs may double the signal. In such an animal, nearly all cilia (20-35 cilia/neuron x millions of OSNs) (43) would be provide a rich source of tissue dedicated to deorphanize that one OR at multiple concentrations with many odors. We are currently characterizing several transgene integration sites in the genome, which will allow expression of any given OR in 50-95% of all dorsal neurons that can offer a streamlined, cost-effective and standardized method to produce mice and define ligands for all human odorant receptors.

### Volatile Odors Emitted by Parkinson’s Disease

In addition to characterizing responses of all human ORs to decode olfaction, our approach can be used to characterize odors emitted by skin or urine, e.g. Comparing the odor profiles of healthy individual over time may reveal odors indicative of a disease state. The use of ORs is different from a general chemical detection platform because it will distill out chemicals or odortopes that are high affinity and have been evolutionarily selected to identify signal from the noise in a complex odor mixture. Recently Gas chromatography-mass spectrometry has been used to identify volatile odors emitted by Parkinson’s Disease (PD) (44). This analysis was born from the surprising observation that a woman, Joy Milne, who happens to be a supersmeller, was able to notice a distinctive “smell” in her husband over a decade before he was diagnosed with Parkinson’s Disease. We envision that a platform with many ORs could distinguish Pre-Parkinson’s Disease odor states from healthy subjects.

## Materials and Methods

### Odorants

Musk compounds were an in-kind contribution from Givaudan. Muscone was purchased from Cayman Chemical, (-)-carvone, and (+)-carvone were purchased from Sigma Aldrich. 3-methyl-2,4-nonanedione was purchased from Santa Cruz Biotechnology and Combi-Blocks.

### Generation of Transgenic Animals

The first, a targeting vector including of IRES-MyrPalm GCaMP6f was generated by swapping *IRES-taumCherry* sequences of the targeting vector, 5×21-OR1A1-IRES-taumCherry, reported in D’Hulst et al (7) with a DNA cassette, *PacI* flanked *IRES-MyrPalm GCaMP6f* with *PacI* site. The 5×21 OR gene choice enhancer sequence (5x ACATAACTTTTTAATGAGTCT) was modified to 9×21 to generate the final targeting vector 9×21-OR1A1-IRES-MyrPalm GCaMP6f. To generate with other OR genes, A *StuI-AscI* fragment which is 187 bases of intron of M71 attached to full CDS of OR1A1 gene was replaced with the *StuI-AscI* fragment with target OR gene CDS (Integrated DNA technologies). Linearized targeting vectors were purified and used for pronucleus injection to obtain founder animals. Founders were bred with CD1 mice. The transgenic mice colonies were maintained by PCR screening for transgene-positive animals. Mice used in this study were bred and maintained in the Laboratory Animal Facility of Hunter College, CUNY. The Hunter College IACUC approved all procedures. Animal care and procedures were in accordance with the Guide for the Care and Use of Laboratory Animals (NHHS Publication No. [NIH] 85-23).

### Characterization of Animals

4 to 12 weeks old mice were used for the analysis by wholemount confocal microscopy imaging with LSM510 (Zeiss). For en face imaging of the GCaMP positive cells, sheets of the olfactory epithelium from the dorsal roof region under the nasal bone were peeled and placed on a slide glass and sealed with live imaging solution (Invitrogen), and coverslipped. Images were taken with UltraView ERS spinning disk confocal microscope (Perkin Elmer). The images were analyzed with FIJI (NIH).

### RT-qPCR analysis

The olfactory epithelia were dissected from 6-week-old mice and stored the tissue at -80°C in RNA*later* Solution (Invitrogen, Cat#: AM7020) until use. Total RNA was extracted using a BEAD BUG6 homogenizer (Benchmark Scientific), the QIAGEN miRNeasy mini kit (Cat#: 217004), and an on-column DNase step (QIAGEN, Cat#: 79254). We used 2.5 μg of total RNA was used to synthesize cDNA via SuperScript IV VILO and we performed the optional ezDNAse step to completely eliminate any DNA contamination (Invitrogen, Cat#: 11766050). qPCR experiments were performed with primers designed via the Primer-BLAST tool on the NCBI website and SYBR-Green (Roche, Cat#: 04707516001) in a Roche LightCycler 480 Instrument II to quantify the results. Each sample/gene pair was loaded in triplicate and each triplicate was repeated across three plates. To calculate Log2 Fold Change, we used a modified ΔΔCt method with corrected primer efficiencies and three reference primers (*Acsm4, Acss2, Slc25a35*).

Additional information and the MiQE checklist and all primer sequences are available in supplemental information. To evaluate the results, we applied t-tests to compare Log2 Fold Change values between genotypes. Due to multiple comparisons, p values were corrected using the Holm-Bonferroni method. Experimental design, calculation of L2FC values, and additional statistical tests are available in supplementary files. R version 4.1.0 was used to convert raw Ct values into L2FC values, perform statistical tests, and generate Fig.S1. Code can be made available upon reasonable request.

### Deciliation of Olfactory Sensory Neurons

Deciliation of OSN was performed with modifications to a previously published protocol (16). Briefly, olfactory epithelium (dorsal recess) from 6-10 week old mice were dissected and placed in ice-cold buffer A (145 mM NaCl, 1.6 mM KCl, 1.6 mM K_2_HPO_4_, 1.2 mM MgSO_4_, 25 mM HEPES, 7.4 mM D-Glucose, cOmplete™ Protease Inhibitor cocktail, pH7.4) then exchanged for ice-cold buffer B (cold buffer A supplemented with 1 mM EDTA, pH7.4) and rocked at 4°C for 15 minutes. The sample is sonicated at low power (two times 20% power for 30 seconds with a 15-second interval) at 4°C, then spun down in a centrifuge at 7,700 x *g* speed for 10 minutes and the supernatant collected in a pre-chilled microcentrifuge tube. The supernatant is spun down at 27,000 x *g* at 4°C for 15 minutes. The resulting pellet is washed once in cold TEM buffer (10 mM Tris, 1 mM EDTA, 3 mM MgCl_2_, pH8.0), then spun down at 27,000 x *g* again for 15 minutes. The pellet is resuspended in 50 μL of TEM buffer with or without 10% glycerol and aliquoted into working volumes, 4-5 μL for subsequent assays. Aliquots are flash-frozen in liquid nitrogen and stored at -80°C until use.

### Ex Vivo Bioassay

Isolated cilia from the dorsal recess are diluted in cold Induction Buffer (25 mM HEPES, 5 mM MgCl_2_, 1 mM ATP, 20 µM GTP, 10 nM GDP, 1 mM DTT) at a dilution of 1:200 from the original TEM+10 % glycerol or without glycerol stock. For liquid delivery of odor, ligands are dissolved in DMSO or ethanol as an intermediate solvent between 50 mM and 1M. Diluted isolated cilia are exposed to ligands pre-dissolved in Induction Buffer for 20 minutes at 37°C. For headspace delivery of odor, ligands are either delivered pure or diluted with dipropylene glycol and placed in an amber vial connected to a custom olfactometer. Odor volume in the vial is set at 100 µL-500 µL, and the dilution is dependent on the calculated saturated vapor pressure delivery amount for a 20 minute odor delivery. Odors delivered pass through a custom 96-well manifold that fits a well plate containing diluted cilia, in which the well plate is heated from below at 37°C. Following this incubation, cAMP is measured using the PerkinElmer AlphaScreen cAMP immunoassay (Cat#6760635) according to manufacturer’s recommendation.

### Data analysis and statistics

The activation of the bioextract of hOR expressing transgenic mice was evaluated by two different calculations. A) activity of each sample by ligands of Tg and WT was calculated by produced cAMP responding to ligands and the solvent (DMSO or Ethanol for liquid delivery, dipropylene glycol for headspace delivery): ΔcAMP = (cAMP^Ligand^-cAMP^solvent^)/cAMP^solvent^ (Fig.3C, 5, 6A, S3, and S5A). B) to evaluate the odors give a negative effect on cAMP production (like Musk compounds), ΔcAMP of each sample (values of A) was corrected with the mean value of it WT to 1.0: ΔcAMP^TG^ + 1 - (mean of ΔcAMP^WT^) (Fig.4, S5B). 3 technical replicates and a minimum of 3 biological replicates. Statistics analyses were performed using Prism 9 (GraphPad).

Mann-Whitney test was performed to evaluate two groups for Fig 3C and 6B. One-way analysis of variance (ANOVA) with Fisher’s least significant difference (LSD) as post hoc comparisons was used to evaluate multiple groups with one independent variable, Fig 5A and 5B. Multiple groups with two independent variables were compared using two-way ANOVA with LSD as post hoc comparison (Fig 4, 6A, S3, and S5).

### S3E and R2 Analysis

Activation of a specific OR is calculated by the Activation values, ΔcAMP at an endpoint measurement. A successful response is defined as a significant (p < 0.05) increase in response of the bioextracts expressing a specific OR when compared to a non-transgenic control sample. We have defined 6 parameters to measure successful profiling of a specific OR (or set of ORs) with a given ligand (or mixture of ligands): Sensitivity, Selectivity, Specificity and Efficacy (S3E) and Repeatability and Reproducibility (R^2^). All 6 parameters can be measured using our existing ex vivo bioassay:

- **Selectivity:** The selected human OR bioextract (n = 3 biological replicates) is stimulated with compounds. A negative control, which non-specifically stimulates the OR, and a positive control, a known ligand specific to the OR, is used in tandem as a benchmark for OR activation. Compounds that generate a response (activation value) significantly higher than the negative control (p < 0.05) are determined as **selective** against the human OR.
- **Specificity:** Ligand response will be compared among wildtype (non-transgenic) and ORs (n = 3 biological replicates). A ligand is identified as **specific** to the OR if the OR differentially responds to the ligand compared to WT or any other OR.
- **Sensitivity:** The specific human OR responds at the lowest concentration to the tested ligand. The ligand that evokes a response at the lowest concentration <u>sets the threshold</u> for the OR and is then identified as the highest-affinity ligand.
- **Efficacy:** A dose-response curve for 4-5 odor concentrations shows a concentration-specific activation profile of the human OR.
- **Repeatability and Reproducibility:** To obtain statistically relevant data, all experiments will be carried out by using 3-5 biological (different animal, same OR) and 3-4 technical (same animal, same OR) replicates per OR and odor concentration.

## Supporting information

Supplemental Statistics File

## Supplementary Materials

- Bioassay_Data.xlsx
- RTqPCR_Data.xlsx
- Statistics.xlsx
- Supplemental Information.xlsx

## Acknowledgments

We are grateful to Givaudan for providing musk compounds. We thank the Hunter College Animal Facility Manager Barbara Wolin and Veterinarian Patricia Glennon for help in maintaining the transgenic colony, and Rada Norinsky and her team of the Transgenic Core Facility at The Rockefeller University for generating transgenic founders. We would like to thank Rosco Technologies/Dr. Dmitry Rinberg for setting up of headspace odor delivery system. Many thanks to Viktoriya Syrovatkina for critical organizational aspects of the project and Nicole Zeak and Mary Slavinsky for Technical support. We thank the rest of the Yesse Technologies team: Elke Giets, Christophe Lemetre, Alvaro Garcia, Raena Mina, Yumi Fujisawa, Luis Hoffman, Chloe Vermierre and Thomas Pauwelyn for their thoughtful contributions to the platform development. A special thanks to Thomas Bozza for review and insightful comments on the manuscript.

## Funding

**NSF SBIR Phase II Award: 1720679; NSF SBIR Phase IIb Award: 1853051. NIH R21 GM126304-01**

## Author contributions

M.O. and P.F. designed constructs and animal analysis experiments; M.O. performed construct cloning and analyzed transgenic animals; Y.T. and S.B. performed QC for ex vivo bioassay; Y.T. and M.O. designed, performed, and analyzed bioassay experiments; E.L. designed, performed, and analyzed qPCR experiments; Y.T., M.O., E.L., and P.F. prepared figures; M.O., P.F., Y.T., C.D., and S.B. wrote the manuscript.

## Competing interests

Previously awarded patent that relates to work: **W02017024028A1**.

## Data and materials availability

If data are in an archive, include the accession number. Mice and Cilia Extracts require an MTA.

## Supplementary Figures

**Fig. S1.**
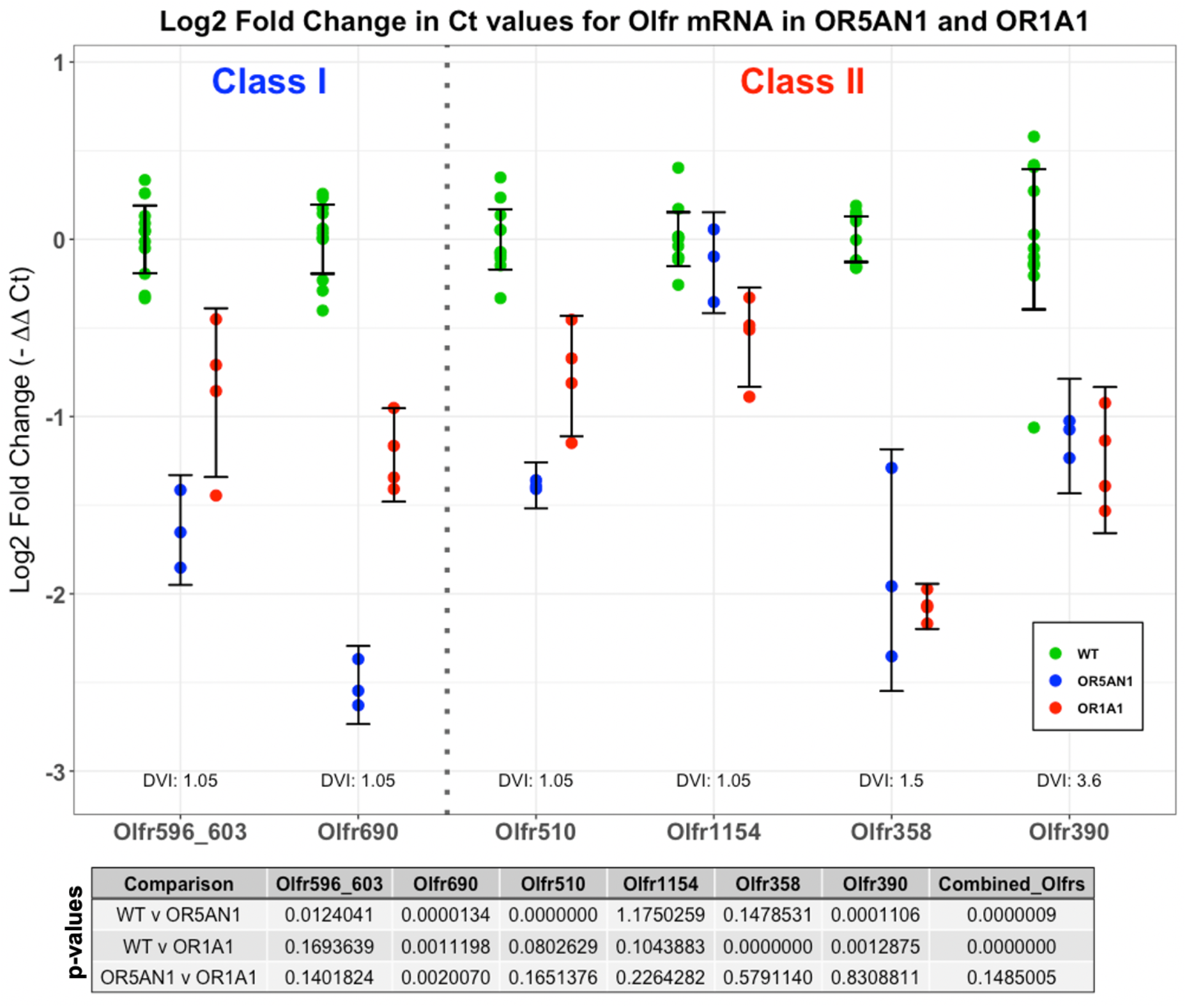
Differential expression of Olfr mRNA in OR1A1 and OR5AN1 transgenic mice. The top panel shows Log2 Fold Change (L2FC) of Olfr mRNA in the olfactory epithelium of OR5AN1 and OR1A1 relative to WT. L2FC calculated using a modified ΔΔCt method to account for primer efficiency and three reference genes (Acsm4, Acss2, Slc25a35). Six Olfr primer pairs were tested, with the Olfr Class and Dorsal/Ventral Index (DVI) displayed. Error bars indicate 95% confidence intervals for each mean. Pairs of Genotypes were compared for significant differences. Testing Olfr genes separately, eighteen t-tests were evaluated, and a Holm-Bonferroni correction was applied for multiple comparisons, the corrected p-values are shown in the table. Combining all Olfr L2FC values for each Genotype, three t-tests were performed, and a separate Holm-Bonferroni correction was applied, the corrected p-values of combined 6 Olfr genes are shown in the most right column. Rounded to seven decimal places, some p values are presented as zero. Biological replicates: WT n = 11, OR5AN1 n = 3, OR1A1 n = 4. Olfr596_603 amplifies both Olfr596 and Olfr603. Alpha = 0.05.

**Fig. S2.**
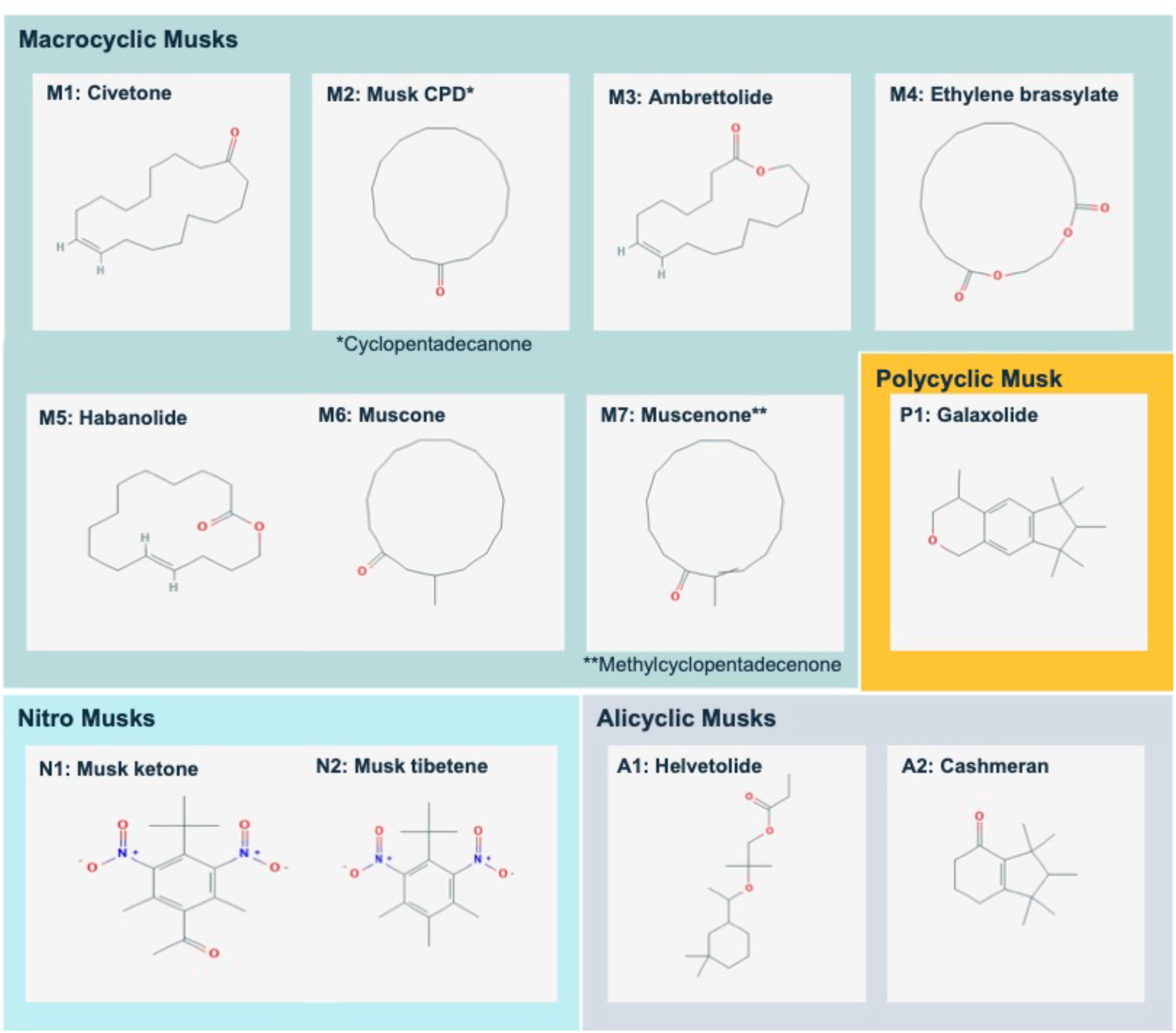
Structure of 12 Musks used for activation of OR5AN1. M = Macrocyclic musks, N = Nitro-musk derivatives, P=Polycyclic musks, A = Alicyclic musks

**Fig. S3.**
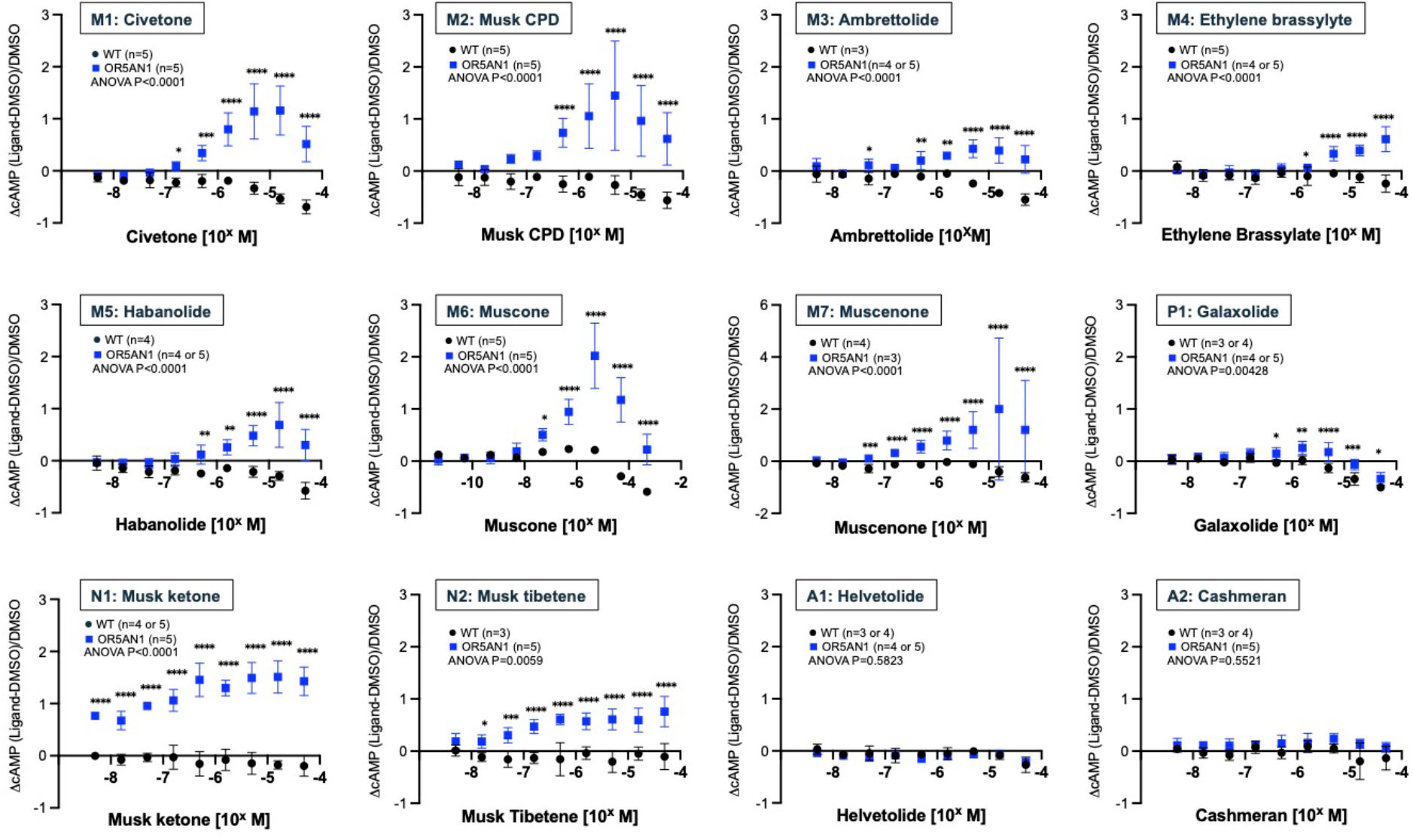
Ex vivo activation of OR5AN1 with 12 musk compounds. Activation of wild type control (WT) and OR5AN1 hOR bioextracts using a nine-point log dilution curve of 12 musk compounds, the black circles and blue squares, WT and OR5AN1 respectively, show mean of, ΔcAMP=(cAMP^ligand^-cAMP^solvent^)/cAMP^solvent^ with ± SD n=3-5 biological replicates as indicated in each graph. Two-way ANOVA was performed, and p-value was as indicated. Fisher’s LSD test was applied as post hoc comparisons, *p<0.05, **p<0.01, ***p<0.001, ****p<0.0001.

**Fig. S4.**
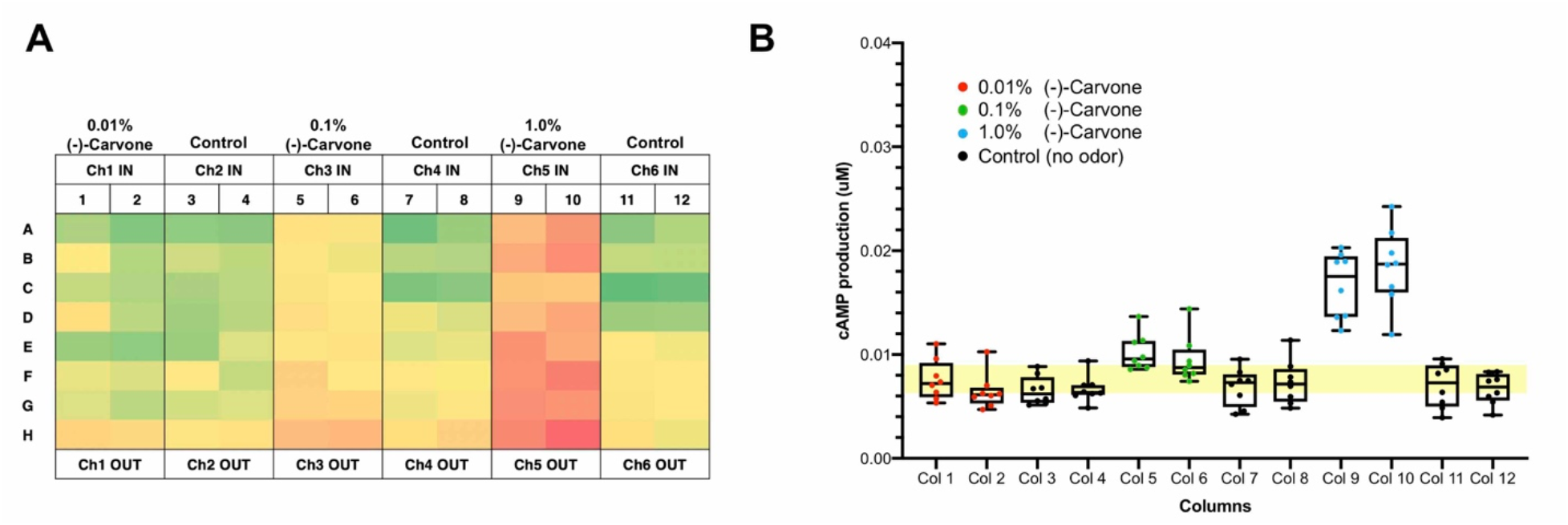
Vapor phase odor delivery system and ex vivo activation of OR1A1 with vapor phase delivered (-)-carvone. Our vapor phase odor delivery system has 6 independent channels to deliver odors to two columns (16 wells) in a 96-well plate at the same time. (**A**) A heatmap of the activation of bioextract of OR1A1 with a series of dilution of (-)- carvone in 96 well plate layout. The level of cAMP production from the wells is shown in green to red color indicator. Diluted (-)-carvones, 0.01%, 0.1% and 1.0% were delivered from channel 1 (Ch1 IN), channel 3 (Ch3 IN), channel 5 (Ch6 IN), respectively. Channel 2, 4, 6 (Ch2 IN, Ch4 IN and Ch6 IN) were control (unopened). (**B**) A nested box plot of cAMP production in OR1A1 bioextract with vapor phase (-)-carvones delivery is shown in (A). Each box show means with minimum to max with SD. The yellow area indicates the mean ± SD of 6 control columns (Col 3,4, 7,8, 11, and 12).

**Fig. S5.**
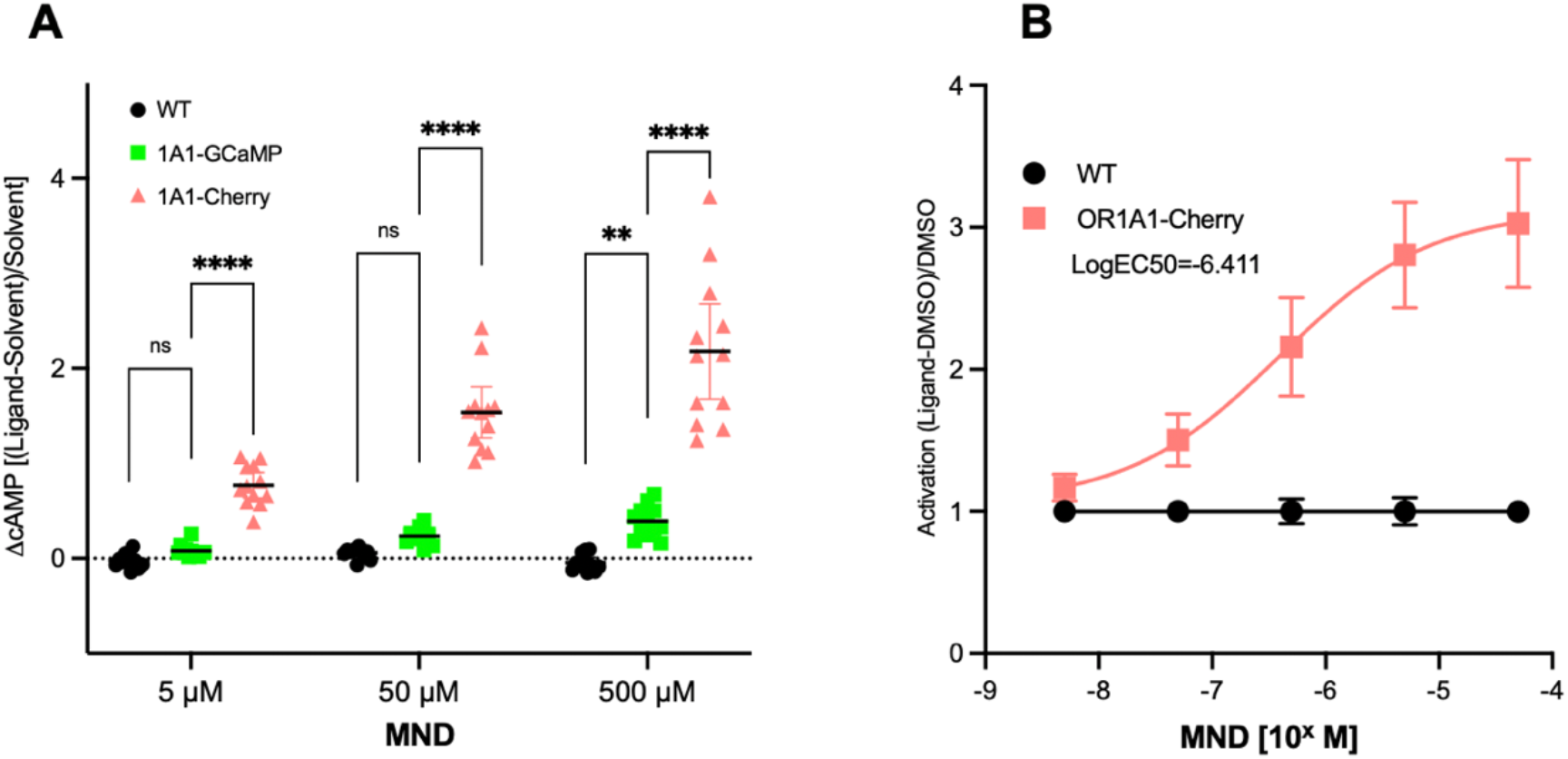
Ex vivo activation of two different OR1A1 transgenic strans with MND. (**A**) The activation of wild type control (WT) and two hOR bioextracts of OR1A1 transgenic strains, 5×21-OR1A1-Cherry (1A1-Cherry) and 9×21-OR1A1-GCaMP6f (1A1-GCaMP) exposed to three concentrations of 3-methyl-2,4-nonanedione (MND). The mean values of, ΔcAMP=(cAMP^ligand^-cAMP^solvent^)/cAMP^solvent^ are shown as black circles, pink triangles and green squares, WT, 1A1-Cherry, and 1A1-GCaMP respectively with SEM. WT; n=11, 1A1-Cherry; n=12, 1A1-GCaMP; n=12. Two-way ANOVA was performed to evaluate ΔcAMP of 1A1-Cherry and 1A1-GCaMP with Fisher’s LSD test was applied as post hoc comparisons, ns at 5μM p=0.459, ns at 50 μM p=0,186, **p= 0.0015, ****p <0.0001. (**B**) Dose-response curves of hOR bioextracts of 5×21-OR1A1-Cherry (1A1-Cherry) with control wild type (WT) to MND. The black circles and pink squares, WT and OR1A1-Cherry respectively, show mean of activation value (ΔcAMP=(cAMP^ligand^-cAMP^solvent^)/cAMP^solvent^) corrected with its WT value=1 ± SEM. WT; n=3, 1A1-Cherry; n=4. LogEC50 was calculated from the best fit nonlinear regression curve (four parameters) with the corrected activation values.

## Notes

### Summary of Updates

Based on reviewer comments we have updated all the figures and included a new figure. In addition, we revised text and made the the data analysis uniform throughout the manuscript.

